# Cell-in-drop genotyping detects megabase-scale loss of heterozygosity as a quantitative marker for CRISPR-Cas9 on-target toxicity

**DOI:** 10.64898/2026.10.30.763197

**Authors:** Darpan Medhi, Linas Mazutis, Maria Jasin

## Abstract

Targeted DNA double strand breaks catalyzed by CRISPR-Cas9 have been demonstrated to cause megabase-scale segmental deletions and whole chromosome losses in a wide variety of mouse and human cell lines. This raises safety concerns for CRISPR-Cas9 based therapeutics and highlights the need for scalable methods to assess its on-target genotoxicity in clinical contexts. In this report, we present cell-in-drop genotyping, as a high throughput approach that combines single cell encapsulation with droplet-based PCR genotyping of endogenous polymorphisms to detect loss of heterozygosity (LOH) at a targeted chromosome without any genome manipulation. With this approach, we detected megabase scale LOH induced by Cas9 DSBs, mostly frequently by segmental loss. This approach also detected LOH from loss of a critical DNA repair factor, BLM helicase, and LOH induced by other genotoxic conditions, in particular, the chemotherapeutic drugs Paclitaxel and Mitomycin C, which were demonstrated to induce LOH by different mechanisms. A modified version of the assay also captured rare local short range LOH arising from DSB-induced interhomologue recombination. Because it requires no genome manipulation, cell-in-drop genotyping offers a practical, scalable tool for evaluating genotoxic risk of CRISPR-Cas9 and other agents in research and clinical development.

## Introduction

Genome editing via CRISPR-Cas9 holds great promise for gene therapy to treat previously incurable diseases [1–13]. However, when it relies on DNA sequence targeted double-strand breaks (DSBs), it has the potential to compromise the genome in unforeseen ways. Concerns regarding the side effects of CRISPR-Cas9 genome editing have largely focused on its off-target activity, reflecting the ability of Cas9 to cleave DNA sequences similar to its target site [14]. High-fidelity variants of Cas9, which have lower tolerance for DNA sequence mismatches, have mitigated this risk to a certain extent, with reduced cleavage of off-target sites [15, 16].

Accumulating evidence has demonstrated that DSBs also produce unwanted repair outcomes at their target sites, *i.e.*, on-target activity, causing the loss of DNA sequences ranging from a few thousand base pairs to millions of base pairs, spanning large chromosome segments and even entire chromosomes [17, 18]. Estimates of these potentially toxic “on-target” events have been reported to be as high as 5-10% in human and mouse cell lines [19–22] and 5-20% in primary human T cells [23, 24]. Other studies have implicated Cas9 DSBs in generating copy-neutral events, such as from reciprocal exchanges between homologous chromosomes at frequencies of ∼0.1-1%, which nonetheless can have profound genetic consequences by leading to loss of genetic information [20, 25–28]. Most dramatically, on-target Cas9 DSBs can also lead to highly rearranged chromothriptic chromosomes [29]. However, despite abundant evidence of on-target genotoxic effects, their occurrence remains largely unaddressed in clinical applications of CRISPR-Cas9 [1–13]. Thus, a critical need exists for clinically applicable methods that can evaluate the on-target toxicity of CRISPR-Cas9 and other DNA damaging gene therapy approaches.

A common outcome underlying many of the megabase scale genomic changes induced by CRISPR-Cas9 is loss of heterozygosity (LOH) across the affected region. LOH is a phenomenon whereby a diploid cell heterozygous for a particular DNA sequence on its two homologues loses one DNA copy and therefore becomes hemizygous in copy loss events or homozygous in copy neutral events. While LOH at tumor suppressor genes is well-known as a critical early event in tumorigenesis [30–33], megabase scale LOH is also relevant to CRISPR-Cas9 genome editing as a consequence of its on-target toxicity [17]. Studies to quantify CRISPR-Cas9’s on-target toxicity have used genome engineering to integrate selectable markers to detect megabase scale LOH [19, 21, 27, 28], but this approach is not a feasible in clinical settings and impractical in many laboratory settings in which genetic manipulation of cells may be difficult.

In this manuscript, we describe a single-cell genotyping assay that detects LOH using existing DNA sequence polymorphisms between chromosome homologues. We refer to our method as cell-in-drop genotyping, in which a large number of cells are individually isolated in water-in-oil droplets and then their genomic DNA is genotyped within an intact diploid context using fluorescent signal from allele-specific PCR-coupled TaqMan probes that correspond to each homologue. Because these reactions are performed by compartmentalizing individual cells, LOH can be detected irrespective of allelic ratios in the overall cell population. This distinguishes our approach from methods that genotype encapsulated purified genomic DNA or genotype cells via barcoding the bulk population followed by cell lysis, both of which may obscure the zygosity of individual cells.

We applied cell-in-drop genotyping to a hybrid mouse embryonic stem cell line which harbors a high density of annotated sequence variations between homologues that facilitate LOH detection. LOH was detected in response to a Cas9-induced DSB as well as genetic and chemical perturbations. By performing cell-in-drop genotyping assays in parallel at both centromere and telomere proximal locations on the edited chromosome, we demonstrated that Cas9-induced megabase deletions mostly resulted from loss of the terminal DNA segment on the edited chromosome. The assay was further adapted to reveal mitotic recombination between two homologous chromosomes via the detection of short range LOH, showing that Cas9 DSB induced inter-homologue homologous recombination is ∼100 fold less frequent than deletions. Thus, cell-in-drop genotyping detects CRISPR-Cas9 induced large genomic events with high sensitivity and can be adapted to assay practically any genomic location with existing DNA sequence polymorphisms.

## Results

### Cell-in-drop genotyping is a high-throughput single-cell method that can detect loss of heterozygosity (LOH) using existing DNA sequence polymorphisms

Given the impact LOH has on the genetic composition of cells, our goal was to devise an approach to detect LOH by genotyping single cells at allelic loci (**Fig. 1A,B**). To develop this approach, we chose a hybrid mouse embryonic stem cell (mESC) line made by crossing 129S1/SvIm (129) and C57BL/6 (B6) mice, which provides an abundance of well-characterized polymorphisms between homologous chromosomes to enable the design of allele-specific primers. Moreover, this cell line also contains tools that can be used to induce LOH [21, 34–36], which will be elaborated upon in later sections. Specifically, we designed allele-specific PCR primers at the *Klf12* gene on chromosome 14 (chr14) using single nucleotide polymorphisms (SNPs) specific to the 129 and B6 genotypes (**Fig. S1A-C**).

**Figure 1:**
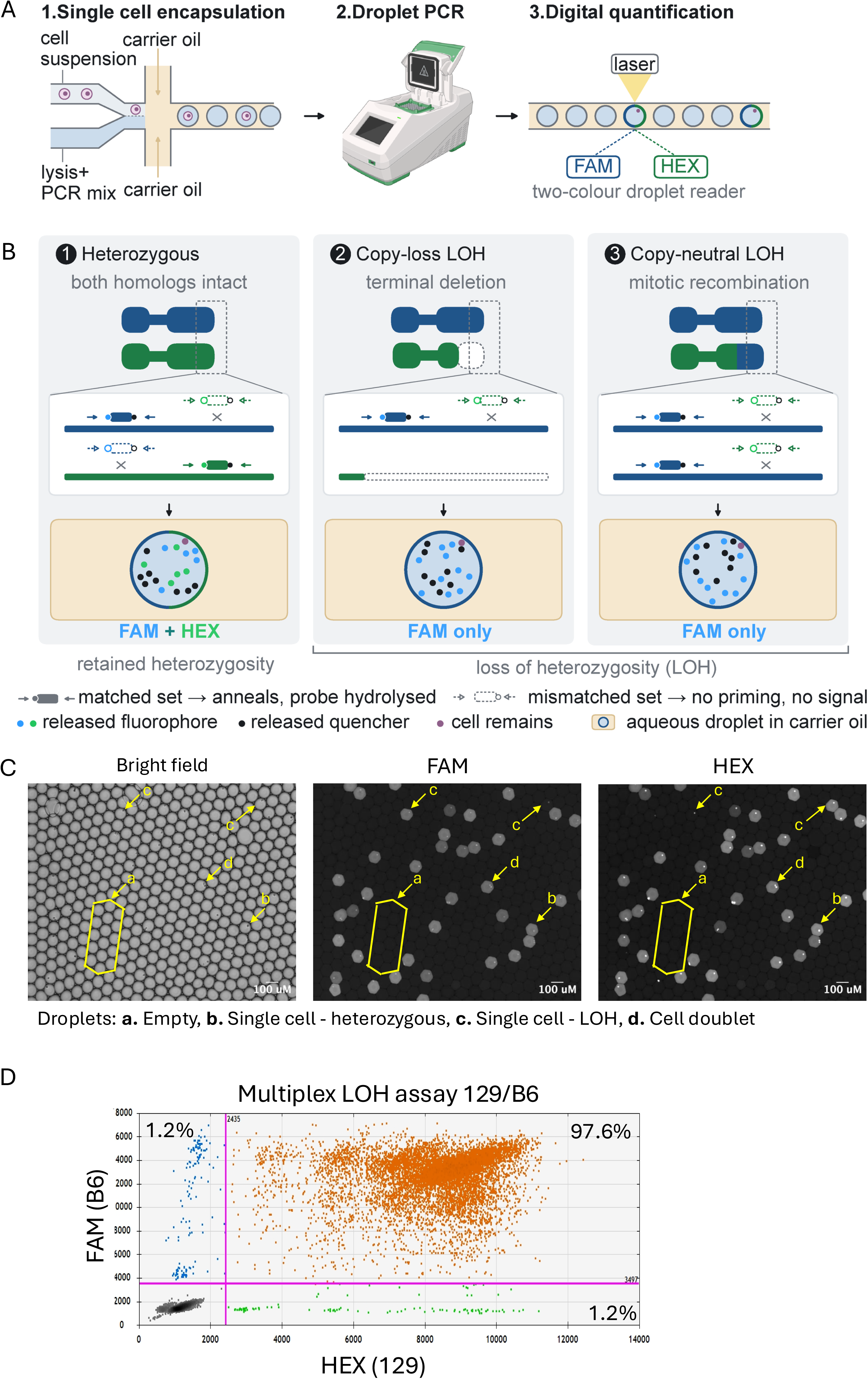
Single-cell genotyping at allelic loci on both homologues via cell-in-drop genotyping detects loss of heterozygosity (LOH). **A.** Schematic of cell-in-drop genotyping workflow. 1. Single cells are encapsulated in water in oil droplets using a microfluidic chip, along with lysis + PCR buffer containing primers and associated fluorescent TaqMan probes. 2. The emulsion is thermocycled post encapsulation, and then 3. The fluorescent signal in droplets is detected with a droplet reader. **B.** Cell-in-drop genotyping assay to detect loss of heterozygosity (LOH). TaqMan probes labelled with fluorophores FAM (blue) and HEX (green) are coupled to allele specific PCR reactions, which are multiplexed to examine different sets of SNPs on each allele. Loss of signal in one channel detects LOH events irrespective of mechanism. Cell remains are the bright foci visible in all PCR signal positive droplets, which are very likely the remaining cell mass after PCR. **C.** Microscopy images showing fluorescence in droplets post PCR. Cells are encapsulated at very low density; hence most droplets are empty and show no fluorescence (a). Amongst droplets with cells, most contain a single-cell and fluoresce in both FAM and HEX (b), indicating heterozygosity. Rare instances of single-cell droplets show fluorescence in only FAM or HEX (c), indicating LOH. Few droplets have >1 cell, often cell doublets (d), and are non-informative for LOH detection but do not affect analysis at their low frequencies. **D.** Post-PCR droplets with cells (as seen in C) can also be analyzed at high throughput using a two-color droplet reader. When allele specific primers were multiplexed to examine 4 different sets of SNPs on each allele, ∼98% of droplets with signal above background were positive for both FAM and HEX, indicating heterozygosity, while ∼1% of droplets each were only single positive for FAM or HEX fluorescence, indicating LOH. The average percent of cells in droplets from all experiments falling within each positive quadrant is indicated.

To isolate individual cells for genotyping, we used the in-Drop platform, which enables ordered high-throughput encapsulation of single cells along with lysis and assay reagents in water-in-oil droplets of controlled volumes [37, 38] (**Fig. 1A**). With this platform, previously developed for single-cell RNA sequencing (scRNA-seq), the cell suspension and reaction buffer are kept separate until cell encapsulation. This prevents premature cell lysis that can generate cell-free DNA fragments, an artifact previously reported in studies using commercial digital droplet PCR (ddPCR) droplet generators to encapsulate cells [39, 40], thereby ensuring that droplets without cells do not show spurious PCR signals (**Fig. 1C**).

For PCR, we utilized Jeffreys’ buffer, which is often used to detect rare sequence variants in cell populations [41]. Using an extended initial denaturation step of 65°C for 1 hour, we found that we could amplify genomic DNA from concentrations ranging from 1 cell in 250 picoliters (pl) reaction buffer to 1 cell in 2 nanoliters (nl) (**Fig. S2A**). We then tested additional components that would be required to effectively perform PCR with cells in droplets, such as non-ionic surfactants that help stabilize such emulsions during thermocycling [42]. We chose non-ionic surfactants whose cloud points (*i.e.,* temperature at which they precipitate from aqueous solutions) are very high (76°C for Tween 20, >100°C for all other detergents) to avoid phase separation at high temperatures and obtained the best DNA amplification results with Tergitol type NP40 and octyl-β-glucoside (**Fig. S2B**). Similarly, we tested different PBS/BSA concentrations and the use of Pluronic F-68 in the cell-suspension buffer, and different concentrations of Mg^2+^ and Taq polymerase in reaction buffers to optimize PCR amplification (**Fig. S2C**).

Having optimized PCR reactions with cells in tubes, we moved onto PCR with cells in droplets and adjusted microfluidic parameters, including droplet volume, the effect of detergent composition on droplet stability and flow rates of various aqueous and oil components. We tested PCR with single cells encapsulated in 500 pl to 1 nl droplets and saw a much lower incidence of PCR failure in droplets below ∼700 pl volume (**Fig. S2D**). For the reaction buffer, we saw the coalescence of fewer droplets post PCR with 0.14% Tergitol type NP40 compared to 0.14% octyl-β-glucoside or no detergent (**Fig. S2E**). With subsequent titration, the Tergitol concentration was increased to 1.4%, which further reduced droplet coalescence. For the final single cell encapsulation conditions, we used flow rates to form ∼700 pl droplets at a rate of ∼500,000 droplets per hour with a density of ∼1 cell per 10 droplets.

TaqMan probes coupled to allele-specific PCR generated fluorescent signal upon genomic DNA amplification. This fluorescence could be directly observed by microscopy (**Fig. 1C**), confirming that ∼10% of droplets contained a single bright focus corresponding to an encapsulated single cell, with a single-cell capture frequency of ≥90%. With an additional increase of the initial denaturation step to 95°C for 20 minutes, a PCR efficiency of ≥99% per cell was achieved in droplets. The single-cell droplets mostly fluoresced in both the B6-targeted FAM and 129-targeted HEX channels, as expected for a population of heterozygous cells, while empty droplets did not fluoresce. There were a few instances of single-cell droplets which only showed fluorescence in the FAM or HEX channel (**Fig. 1C**). These were droplets containing cells that either had undergone LOH at the locus prior to amplification or in which one allele failed to amplify. A small proportion of (<1%) droplets had more than one cell (indicated by >1 bright focus), which would score by fluorescence as heterozygous, but at frequencies too low to substantially affect LOH quantification. Post PCR droplets were read at high throughput with a two-color droplet reader (Bio-Rad QX200) (**Fig. 1D and Fig. S2F**).

We initially used a duplex PCR approach, with one set of primers and probes for each allele (**Fig. 1B)**, and, as a control, encapsulated an equal mix of cells that were homozygous for either the 129 or B6 alleles. Nearly 98% of the droplets were single-positive for either FAM or HEX, demonstrating high sensitivity for LOH detection (**Fig. S2F**). About 2% of droplets were double-positive, likely due to encapsulated cell doublets. However, when 129/B6 heterozygous cells were encapsulated, we found that only ∼88% of the cell-droplets were double positive, with an unexpectedly high level of single-positives (∼12%) (**Fig. S2F**), likely due to allele dropout from PCR failure.

To overcome this, we multiplexed the PCR approach to simultaneously detect four sets of SNPs for each allele (**Fig. S1B,C**). With this approach, 97.6% of the cell population were double-positive (**Fig. 1D**). Thus, multiplexing greatly reduced allelic dropout and improved the sensitivity of the method. About 2.4% of the population remained single-positive for either FAM or HEX (**Fig. 1D**), which could reflect true LOH events, residual allelic dropout, or a mixture of both. We then proceeded to validate our LOH detection using biologically relevant perturbations with tools we established in our prior studies [21, 36].

### Validation of LOH detection by cell-in-drop genotyping

In addition to heterozygosity between homologous chromosomes, the 129/B6 hybrid mESC line cell line contains doxycycline repressible alleles of the BLM helicase [43], a crucial protein for maintaining genomic stability. Also present in these cells is a cleavage site for the I-SceI homing endonuclease on the 129 chr 14, within the *S2neo* gene inserted at the *Rb1* locus, located ∼26.5 Mb centromere proximal to *Klf12* (**Fig. 2A**). Loss of the 129 *Klf12* polymorphisms upon I-SceI cleavage, therefore, indicates extensive LOH from a DSB. These conditions – DSB formation and BLM depletion – perturb LOH by distinct mechanisms: DSB formation is directed specifically to a single chromosome homologue, while BLM depletion has a genome-wide impact.

**Figure 2:**
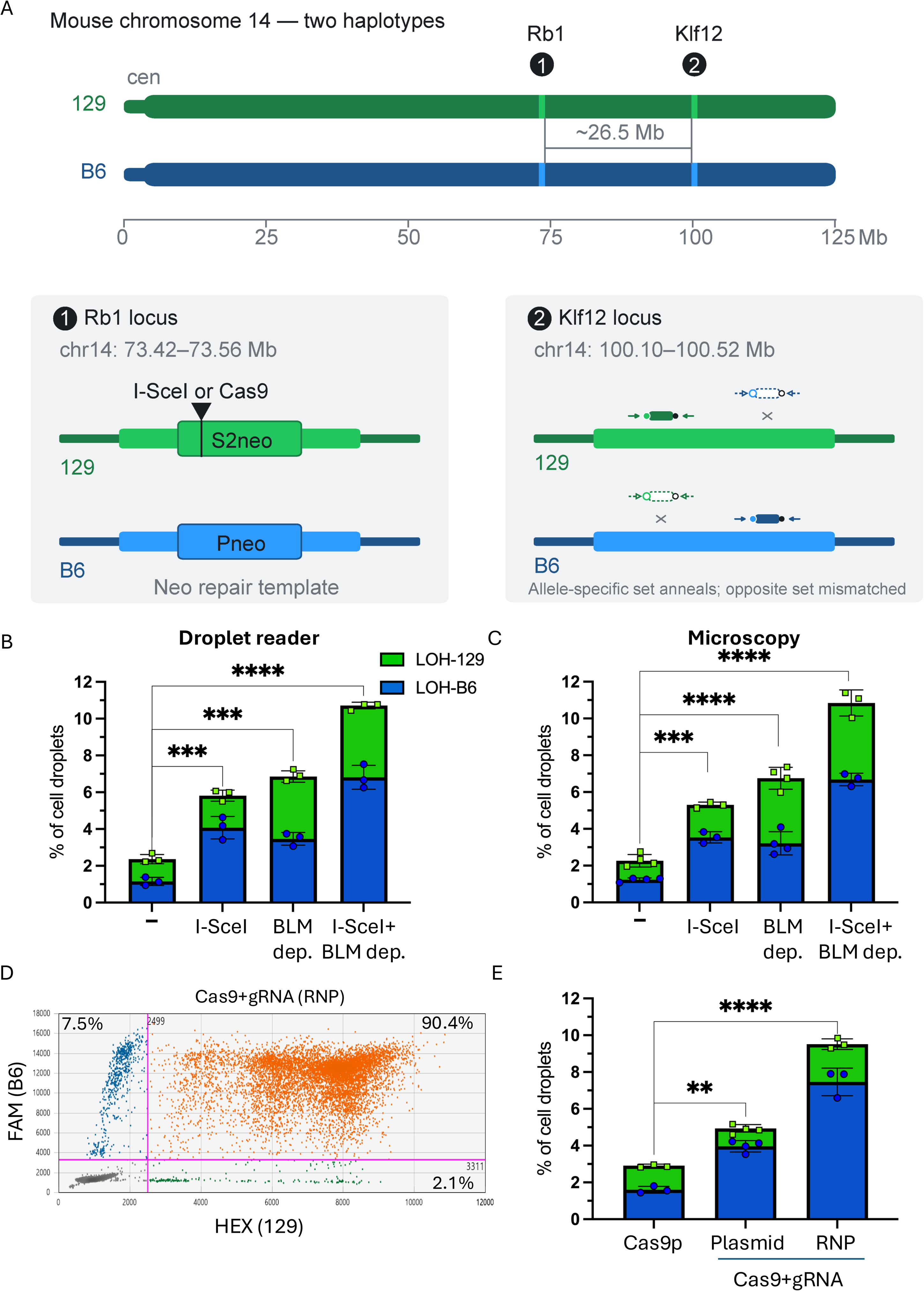
Cell-in-drop genotyping detects increased LOH levels in response to biologically relevant perturbations, including DSBs induced by Cas9. **A.** DSB sites are introduced within the *Rb1* gene on chr14, which is 26.5 Mb from the *Klf12* locus to which allele specific Taqman primers and probes are designed. I-SceI and Cas9 endonuclease sites are present on the 129 chromosome *S2neo* gene embedded within *Rb1*. The *Pneo* repair template is also embedded within *Rb1* but on the B6 chromosome. **B.** Quantification of cell droplets single positive for either FAM (blue, LOH-B6) or HEX (green, LOH-129) under the indicated conditions using a two-color droplet reader. I-SceI leads to a preferential increase in LOH-B6 due to DSB-induced loss of 129 chromosomal sequences, whereas BLM depletion impacts the 129 and B6 chromosomes similarly. >10,000 cells were counted for each condition and significance was determined with ordinary one-way ANOVA. **C.** Quantification of cell droplets single positive for either FAM (LOH-B6) or HEX (LOH-129) under the indicated conditions manually inferred from fluorescence microscopy images. >1000 cells were counted for each condition and significance was determined with ordinary one-way ANOVA. **D.** High throughput analysis for LOH at *Klf12* with the droplet reader in 129/B6 cells after transfection of Cas9 and gRNA RNP. The average percent of cells in droplets from all experiments falling within each positive quadrant is indicated. **E.** Quantification of single positive cell droplets as in **B**. Cas9 cleavage leads to a preferential increase in LOH-B6 due to DSB-induced loss of 129 chromosomal sequences, with RNP transfection being more effective at DSB introduction than plasmid transfection.

To examine the impact of single or dual perturbation using cell-in-drop genotyping, the 129/B6 mESCs were transfected with a plasmid expressing I-SceI endonuclease with or without prior doxycycline treatment to deplete BLM. An increase in single positive droplets was observed under all three conditions, indicating elevated LOH levels (**Fig. 2B and Fig. S3A-C**). After I-SceI expression, the fraction of single FAM positive cell droplets increased by ∼3%, while the number of single HEX-positive droplets increased by ∼0.5%; thus, overall LOH levels increased by ∼3.5% (p=0.0009) (**Fig. 2B and Fig. S3A**). DNA damage specifically directed to the 129 chr 14 therefore led to a specific increase in cell droplets retaining only the B6 allele, termed LOH-B6. On the other hand, BLM-depleted mESCs also showed an increase in single positive droplets (∼4.5%, p=0.0001) (**Fig. 2B and Fig. S3B**); in this case the increase was ∼2.2% for LOH-129 (cell-droplets retaining only the 129 allele) and ∼2.3% for LOH-B6, *i.e.,* roughly similar loss for each of the alleles. The symmetrical LOH is consistent with BLM depletion having an equal chance of affecting either homologue due to global, spontaneous genomic instability. Combining I-SceI expression and BLM depletion led to a roughly additive increase in LOH (**Fig. 2B and Fig. S3C**), suggesting that LOH induced by these two perturbations arises via two distinct mechanisms.

The above cell-in-drop genotyping results from the I-SceI endonuclease and BLM perturbations were further independently verified by examining post-PCR droplets via fluorescence microscopy (**Fig. S4A-D**). LOH levels determined in this way (**Fig. 2C**) were similar to those estimated via the high-throughput reader, further validating use of the two-color droplet reader for high-throughput analysis.

### Cell-in-drop genotyping detects extensive megabase-scale LOH induced by Cas9 DSBs

Many therapeutics under development for clinical applications utilize CRISPR-Cas9 DSBs for efficient genome modification [44]. However, given its potential for generating megabase-scale LOH, we asked whether cell-in-drop genotyping could be used to detect Cas9-induced LOH. We generated a Cas9 DSB with a gRNA targeting the *S2neo* gene on the 129 chromosome adjacent to the I-SceI cleavage site [21] (**Fig. 2A**). Transfecting either a plasmid expressing Cas9 and the gRNA or a ribonucleoprotein (RNP) complex of Cas9 protein and the gRNA substantially increased LOH-B6 compared to mock transfection of Cas9 protein alone (Cas9p; **Fig. 2D,E and Fig. S3D,E**). Cells transfected with plasmid DNA showed increased LOH (∼2%, p=0.0031) entirely driven by an increase in LOH-B6 (**Fig. 2E and Fig. S3E**). RNP transfection led to an even greater increase in LOH (∼6.6%, p<0.0001), again almost all driven by increased LOH-B6, presumably reflecting the more efficient DSB formation by RNP compared to plasmid expression (**Fig. 2D,E**). Thus, cleavage of the 129 chromosome by either Cas9 or I-SceI led to LOH retaining the uncut B6 chromosome, indicating the mode of enzymatic cleavage, *i.e.,* leading to blunt and 3’ overhangs, respectively, had no discernible impact on LOH. These experiments demonstrate the potential for cell-in-drop genotyping to quantify the on-target genotoxic effects of CRISPR-Cas9 genome editing using existing DNA polymorphisms, an approach that could find utility in clinical settings as well.

### Adapting cell-in-drop genotyping to detect short range LOH from IH-HR

Mechanistically, extensive LOH arising from a DSB has been most often associated with large segmental deletions or even whole chromosome loss [19–21, 23, 24, 28, 45–47]. However, some studies have claimed that minimal LOH induced by Cas9 DSBs in embryos can arise from frequent interhomologue homologous recombination (IH-HR) [48, 49] and proposed Cas9 DSB-induced IH-HR as a genome editing strategy that does not require exogenous DNA. The reported frequencies of IH-HR in these studies were ∼1000-fold higher than IH-HR frequencies previously determined in mESCs [35, 36]. However, since the latter frequencies were determined by colony forming assays requiring drug selection, the possibility remained that IH-HR frequencies are higher in cells than previously estimated.

We adapted cell-in-drop genotyping to directly detect short range LOH arising from IH-HR, whereby a DSB site on one homologue is repaired by copying sequences from the uncut homologue (**Fig. 3A** and **Fig. S5A**). Allele-specific primers were designed to the mutant *S2neo* and *Pneo* genes on the two homologues, such that when used together they would only amplify the *neo^R^* gene arising from IH-HR, which is detectable with a HEX probe (**Fig. S5B**). We combined the IH-HR primers with a set of universal control primers upstream of the *neo* genes to identify all cells using a FAM probe (**Fig. S5B**). We tested this duplex IH-HR assay with a population of unperturbed mESCs (no DSB) and a population of previously isolated *neo^R^* cells. Amplification from the allele specific primers from the unperturbed cells was very low (∼0.003-0.006%); conversely, most *neo^R^* cells were positive for both FAM and HEX (∼82%) **(Fig. S6A,B)**. Another ∼7% of *neo^R^* cells were only HEX positive. Since the HEX signal in these droplets was comparable to the HEX signal in double positive droplets, they likely represent *neo^R^* cells with allelic dropout of the FAM PCR reaction.

**Figure 3:**
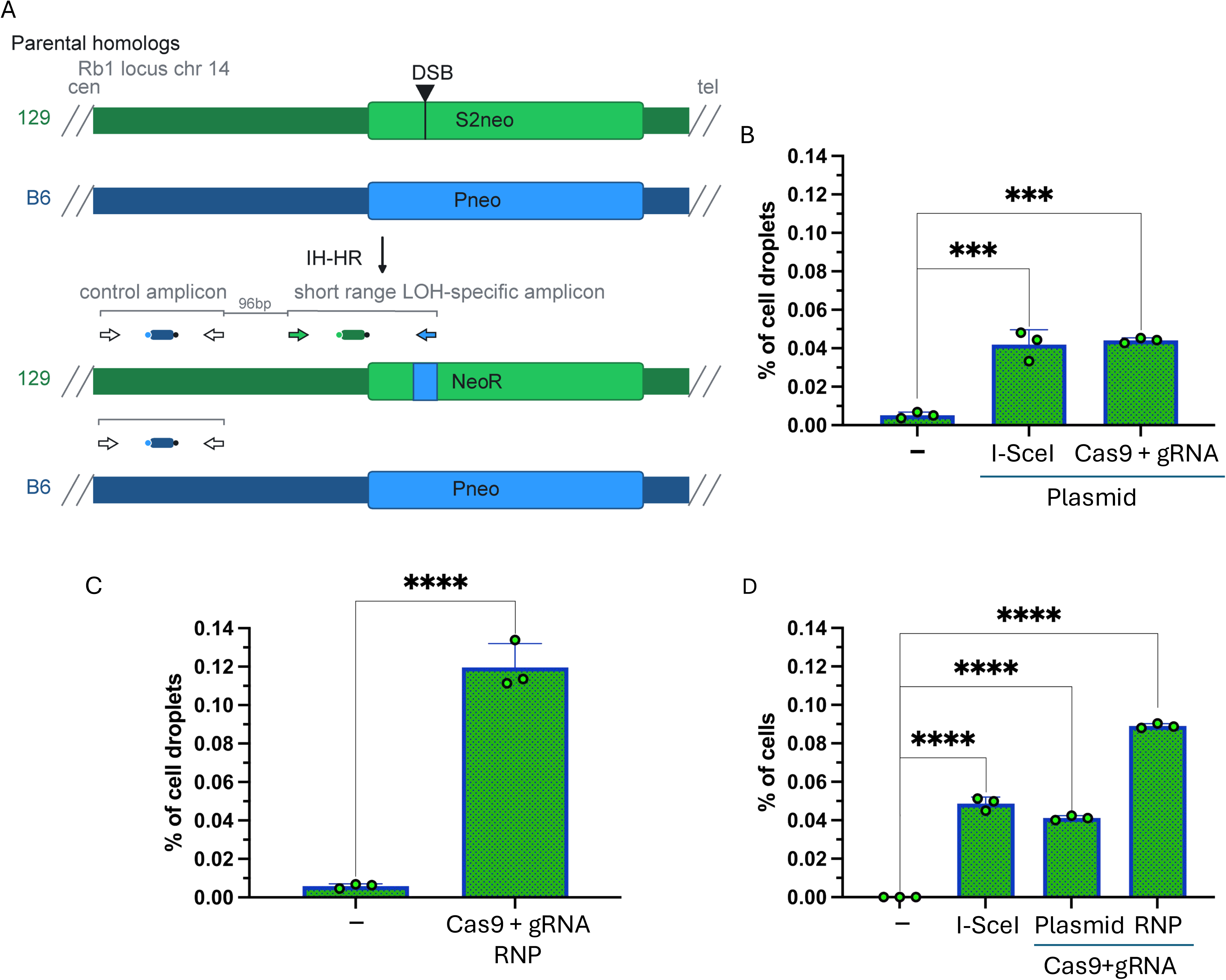
Cell-in-drop genotyping detects short range LOH from inter-homologue homologous recombination (IH-HR). **A.** IH-HR system in 129/B6 in mESCs. A DSB at the *S2neo* locus repaired from homologous sequences on the *Pneo* locus results in a *neo^R^* gene with short-range LOH (blue box in the *neo^R^* gene). A HEX-labelled TaqMan probe identifies the IH-HR specific PCR product generated with the forward primer on the 129 allele and the reverse primer on the segment with LOH retaining B6 sequences. TaqMan probes labelled with FAM are coupled to universal (non-allele specific) control PCR reactions upstream of the *neo* genes. In the multiplex approach, two control primer sets are used (not shown), whereas in the duplex approach, only one primer set is used (as shown). **B.** Quantification of IH-HR levels for all tested conditions from the duplex IH-HR assay using a two-color droplet reader. >30,000 cells were counted for each condition, and significance was determined by ordinary one-way ANOVA. **C.** Quantification of IH-HR levels for both tested conditions from the multiplexed IH-HR assay. >30,000 cells were counted for each condition, and significance was determined by ordinary one-way ANOVA. **D.** Quantification of IH-HR levels determined via *neo^R^* colony formation. Significance was determined with ordinary one-way ANOVA.

We then induced IH-HR with a DSB targeted to the *S2neo* gene using plasmids expressing either I-SceI or Cas9 and performed cell-in-drop genotyping two days later. FAM and HEX double positive and HEX single positive cell droplets comprised ∼0.04-0.05% of the cells, about 10-fold above background (**Fig. 3B and Fig. S6C,D**), demonstrating that even relatively rare IH-HR events can be detected by cell-in-drop genotyping.

To reduce allelic dropout from the FAM channel, we multiplexed the assay to include a second control FAM reaction and used a Cas9 RNP to induce a higher level of cleavage at the *S2neo* locus. FAM and HEX double positive droplets increased to ∼0.12%, while the level in the absence of a DSB was unchanged at 0.005% (**Fig. 3C and Fig. S6E,F**). With multiplexing, the proportion of single-HEX-positive droplets was reduced compared with double-positive droplets, consistent with reduced allelic dropout.

We compared cell-in-drop genotyping with the previous genetic assay for IH-HR which involved selection for *neo^R^* colonies ∼10 days after DSB induction [21, 35, 36] (**Fig. S6G)**. Similar frequencies of *neo^R^*colonies were observed as with cell-in-drop genotyping (**Fig. 3D**). These experiments demonstrate that short range LOH from IH-HR can be readily detected by cell-in-drop genotyping, but it is infrequent relative to the long range LOH observed after a DSB.

### Insight into Cas9-driven LOH mechanisms through parallel cell-in-drop genotyping at centromere and telomere proximal loci

Our cell-in-drop genotyping results are consistent with prior studies that a single Cas9 DSB leads to LOH typically through mechanisms other than IH-HR [17, 18]. In particular, using a fluorescence-based system, we previously observed frequent segmental loss, especially from chromosome truncations, and less frequently whole chromosome loss [21]. Assessing LOH near the ends of the chromosome differentiates between these two mechanisms, as cells undergoing whole chromosome loss lose heterozygosity at both ends, while cells with segmental loss retain heterozygosity near the centromere. Thus, at a population level with whole chromosome loss, LOH at centromere and telomere proximal loci would occur at similar levels, while with segmental loss, LOH would be more frequent near the telomere than near the centromere (**Fig. S7**).

To address this, we used allele-specific PCR with TaqMan probes to the *Fhit* locus located ∼11 Mb from the centromere and to the *Klf12* locus located ∼25 Mb from the distal telomere (**Fig. 4A and Fig. S8A-C**). A DSB was targeted 0.8 Mb centromere proximal to the *Klf12* locus specifically on the B6 chromosome and then cell-in-drop genotyping was performed in parallel at both the *Fhit* and *Klf12* loci (**Fig. S7**). LOH-129 at *Klf12* increased substantially to 5.8% of the population with the Cas9 RNP (p<0.0001), while LOH-129 at *Fhit* only increased by only 0.5% (p=0.2584) (**Fig. 4B and Fig. S9A,B**). By contrast, LOH-B6 was minimally affected at either locus (<0.5%). These results imply that Cas9 DSB induced LOH events are primarily driven by megabase-scale segmental loss from the cleaved chromosome end rather than whole chromosome loss, and that cell-in-drop genotyping can readily quantify such events in a population (**Fig. 4B**).

**Figure 4:**
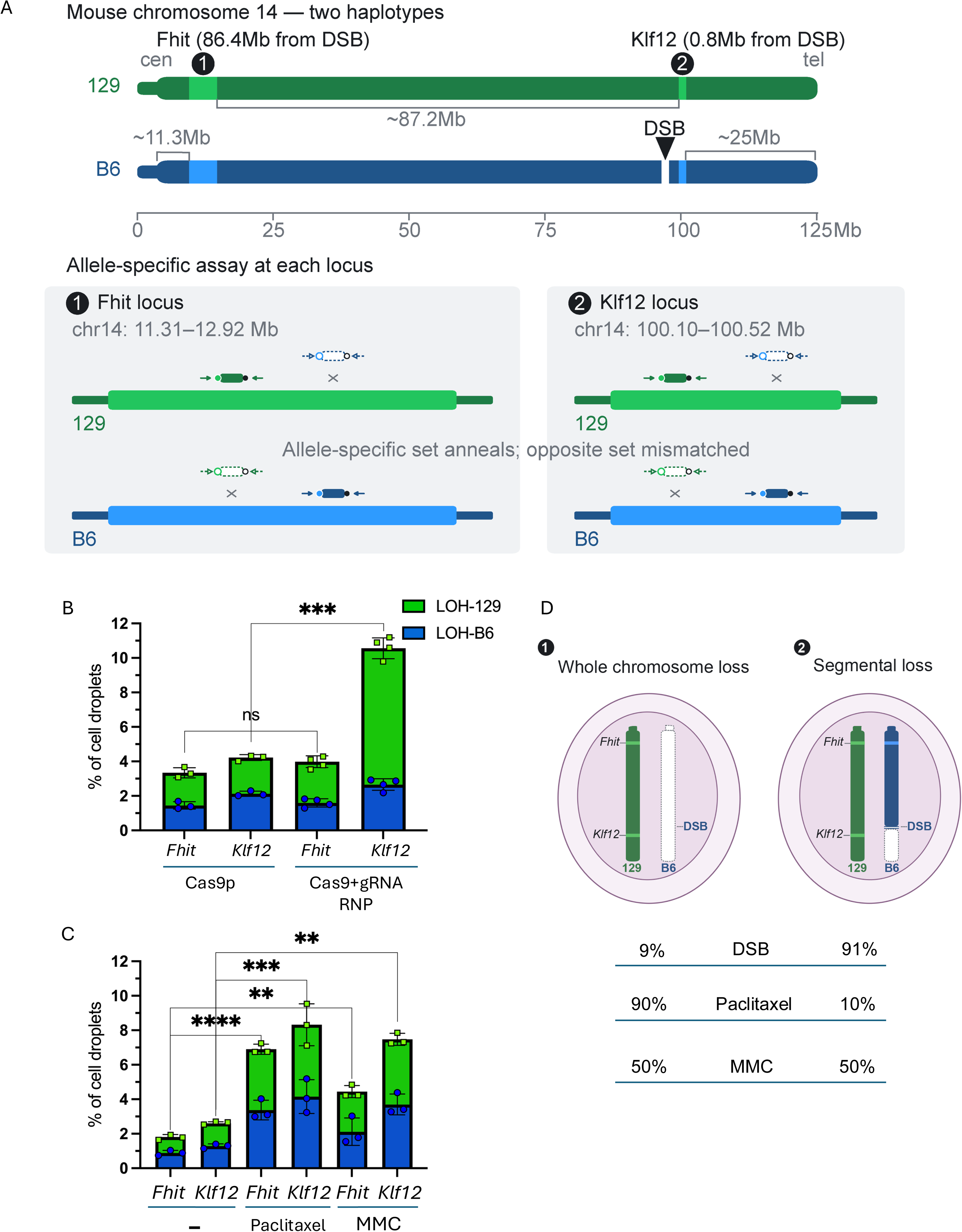
Parallel cell-in-drop genotyping provides mechanistic characterization of LOH by demonstrating that Cas9 DSB induced LOH is primarily driven by segmental loss, while Paclitaxel and Mitomycin C treatments lead to distinct LOH patterns consistent with their genotoxic effects. **A.** TaqMan probes labelled with FAM and HEX are respectively coupled to 129 and B6 allele-specific primers located at the *Fhit* locus ∼11Mb from the centromere, and at the *Klf12* locus ∼25Mb from the telomere for parallel allele specific assays at each locus. **B.** Quantification of cell droplets single positive for either FAM (LOH-B6) or HEX (LOH-129) using a two-color droplet reader in response to transfection of Cas9 protein alone or Cas9 RNP complex inducing a DSB ∼0.8 Mb upstream of *Klf12*. >10,000 cells were counted for each condition, and significance was determined with ordinary one-way ANOVA. **C.** Quantification of cell droplets single positive for either FAM (LOH-B6) or HEX (LOH-129) in untreated cells or cells treated with Paclitaxel or Mitomycin C using a two-color droplet reader. >10,000 cells were counted for each condition, and significance was determined with ordinary one-way ANOVA. **D.** LOH increase driven by a telomere proximal DSB is almost all from the 129-*Klf12* locus, consistent with LOH being primarily from segmental loss on the B6 chromosome. Paclitaxel treatment leads to a comparable increase in LOH at both *Fhit* and *Klf12* loci, consistent with LOH derived primarily from whole chromosome loss (WCL). Finally, Mitomycin C treatment leads to an increase in LOH at both *Fhit* and *Klf12* loci, but the increase at *Klf12* is almost double, indicating that both whole chromosome loss and segmental loss drive LOH. The estimates for LOH are derived by subtracting the background level of single positive droplets from that of either DSB induced, Paclitaxel treated or Mitomycin C treated cells and then dividing the total *Fhit* single positives by the total *Klf12* single positives to derive the approximate percent of WCL events.

### Distinct LOH patterns from chemotherapeutics Paclitaxel and Mitomycin C

Genome destabilizing agents used in cancer therapy also have the potential to cause LOH. We tested two chemotherapeutics which differentially impact the genome, the microtube-stabilizing drug Paclitaxel [50] and the DNA cross-linking agent Mitomycin C [51, 52]. mESCs were treated with sublethal doses of Paclitaxel (3.25 nM) or Mitomycin C (15.63 nM) for 72 hours and then analyzed by cell-in-drop genotyping together with untreated cells (**Fig S9C-E**). At these doses, both agents led to significantly increased levels of single-positive cell droplets in both the FAM and HEX channels (**Fig. 4D**), consistent with the lack of specificity of either agent for a particular chromosome. Compared to no treatment, Paclitaxel treatment led to a similar net increase in single positive FAM and HEX droplets at both *Fhit* and *Klf12* (5.1% and 5.7%, respectively; p=0.9690) (**Fig. 4D and Fig. S9 C,D**). Conversely, Mitomycin C treatment led to a greater increase in single positive cell droplets at *Klf12* than at *Fhit* (4.9% and 2.6%, respectively; p=0.0244) (**Fig. 4C and Fig. S9C,E**). These observations are consistent with Paclitaxel-induced LOH being primarily driven by whole chromosome loss, while Mitomycin C induced LOH involves a mixture of whole chromosome and segmental loss (**Fig. 4D**).

## Discussion

In this work, we developed cell-in-drop genotyping as a high-throughput targeted approach for detecting chromosome-specific LOH at single-cell resolution. Cell-in-drop genotyping detected megabase-long LOH induced by either a DSB or depletion of a critical DNA repair factor (BLM), or a combination of both, finding LOH in up to 10% of cells, similar to what was observed with cells engineered to contain an integrated fluorescent LOH reporter [21]. In addition to megabase-scale LOH, we also adapted the approach to detect highly local short-range LOH arising from DSB-induced interhomologue recombination, which was much less frequent. Parallel LOH assays with probes to centromere and telomere proximal loci demonstrated DSB-induced LOH to be largely driven by segmental loss, as would be expected from chromosome truncations [21]. Extending the analysis to cells treated with sublethal doses of chemotherapeutics, we observed that Paclitaxel primarily leads to LOH driven by whole chromosome loss, consistent with disrupted mitotic chromosome segregation [50], while Mitomycin C leads to LOH driven by a combination of whole chromosome and segmental loss, in line with LOH driven by a random distribution of DNA crosslinks along the chromosome [51, 52]. LOH detection with cell-in-drop genotyping upon chemotherapeutic treatment provides an important, novel approach to measuring genotoxicity, because in addition to determining LOH levels, the mode of LOH can be discerned at high throughput, *i.e.,* whole chromosome versus segmental loss, providing insight into mechanisms of action. LOH detection in all cases was possible with cell-in-drop genotyping because template DNA is partitioned through individual cell separation, such that genotyping occurs in an intact cellular and genomic context, unlike other digital PCR methods.

As with other LOH detection methods, results from cell-in-drop genotyping raise safety concerns for CRISPR-Cas9 genome editing therapeutics. While the presence of chromosomal aberrations leading to LOH is not necessarily an indication of an adverse clinical event, genomic instability is a recognized hallmark of tumorigenesis, implying a need for further investigation and possible intervention. However, of the 79 clinical trials currently listed on the clinicaltrials.gov website that utilize CRISPR-Cas9 genome editing of which 33 are active, methods to detect on-target chromosome instability have not been disclosed in any of them [53]. In the studies that do provide this information, analyses involved a one-time characterization of edited cells, with no long-term patient surveillance. Patient surveillance in these studies primarily focused on malignancies, which is a lagging indicator where the harm to the patient has already occurred. Moreover, chromosome instability monitoring, when provided, appears to have overlooked infrequent megabase-scale losses. For example, in the European assessment report (EMA/6332/2024) for the BCL11A-enhancer clinical trials for beta-thalassemia and sickle cell anemia (NCT03655678 and NCT03745287), limited karyotyping (60 metaphases) and sequencing around the DSB site (∼10 kb) was performed. However, Cas9 DSBs frequently result in loss of centromere proximal DNA sequences extending several megabase pairs, such that localized analyses around the DSB site will overlook these events [21]. More recent studies have utilized methods such as CAST-Seq, which has a similar limitation of only detecting events in the direct vicinity of the DSB site, or scRNA-seq, which relies on large gene sets and thus may overlook segmental loss induced, for example, by telomere proximal Cas9 DSBs [24, 54, 55].

To overcome the limitations of existing approaches, we propose cell-in-drop genotyping as a targeted approach to monitor CRISPR-Cas9 on-target toxicity, which detects megabase-scale losses with high sensitivity and works without a requirement for prior genome engineering or cell selection, unlike many other methods used in cellular studies [21, 22, 28, 29]. The main prerequisite for our approach is knowledge of SNPs that enable the design of allele specific PCR reactions. While this information has been historically lacking, increasing numbers of annotated SNPs from the NCBI dbSNP database or *de novo* annotated and haplotyped SNP maps of human cells can simplify the design of allele specific PCRs. In addition to cell-in-drop genotyping, targeted single-cell DNA sequencing (scDNA seq) has recently been developed to detect CRISPR-Cas9 induced megabase-scale LOH [56]. This method used ∼400 SNPs distributed on two human chromosomes for LOH detection, while cell-in-drop genotyping in principle only requires ∼8 SNPs near the telomere to detect LOH on any chromosome. Cell-in-drop genotyping has other potential advantages, for example, rapid fluorescent signal readout, rather than bioinformatic analysis of sequencing data, and the ability to detect low levels of background LOH. Finally, cell-in-drop genotyping can be combined with droplet sorting based on the fluorescence from TaqMan probes, which could then enable subsequent analysis of the encapsulated genomes, transcriptomes and proteomes [57].

## Acknowledgements

We thank members of the Jasin lab, for discussions and support, especially Jung Eun Kim and Travis White for comments on the manuscript. We are also grateful to Andrea Farina and members of the MSKCC Integrated Genomics Operation (IGO), for technical support and sharing of resources, especially for high throughput analysis with digital droplet PCR readers. This work was supported by MSKCC Basic Research Innovation fellowship 239218, Fund 16621 (D.M.), Cancer Center Core Grant P30CA008748 and R35CA253174 (M.J.).

## Author contributions

D.M., L.M., and M.J. conceived the study; D.M. performed the experimental work; D.M., L.M., and M.J. wrote the manuscript.

## Policy and Ethical Declarations

The authors declare that the AI tool Claude Science was utilized to generate initial conceptual templates for the explanatory graphics in Figures 1A, B; 2A; 3A; 4A, D; S1A, S5A, S6G, S7 and S8A. These templates were then extensively reviewed, verified, and edited by the authors to guarantee scientific accuracy and clarity. The authors assume full responsibility for the integrity of the final visual content.

## Supplementary figure legends

**Figure S1:**
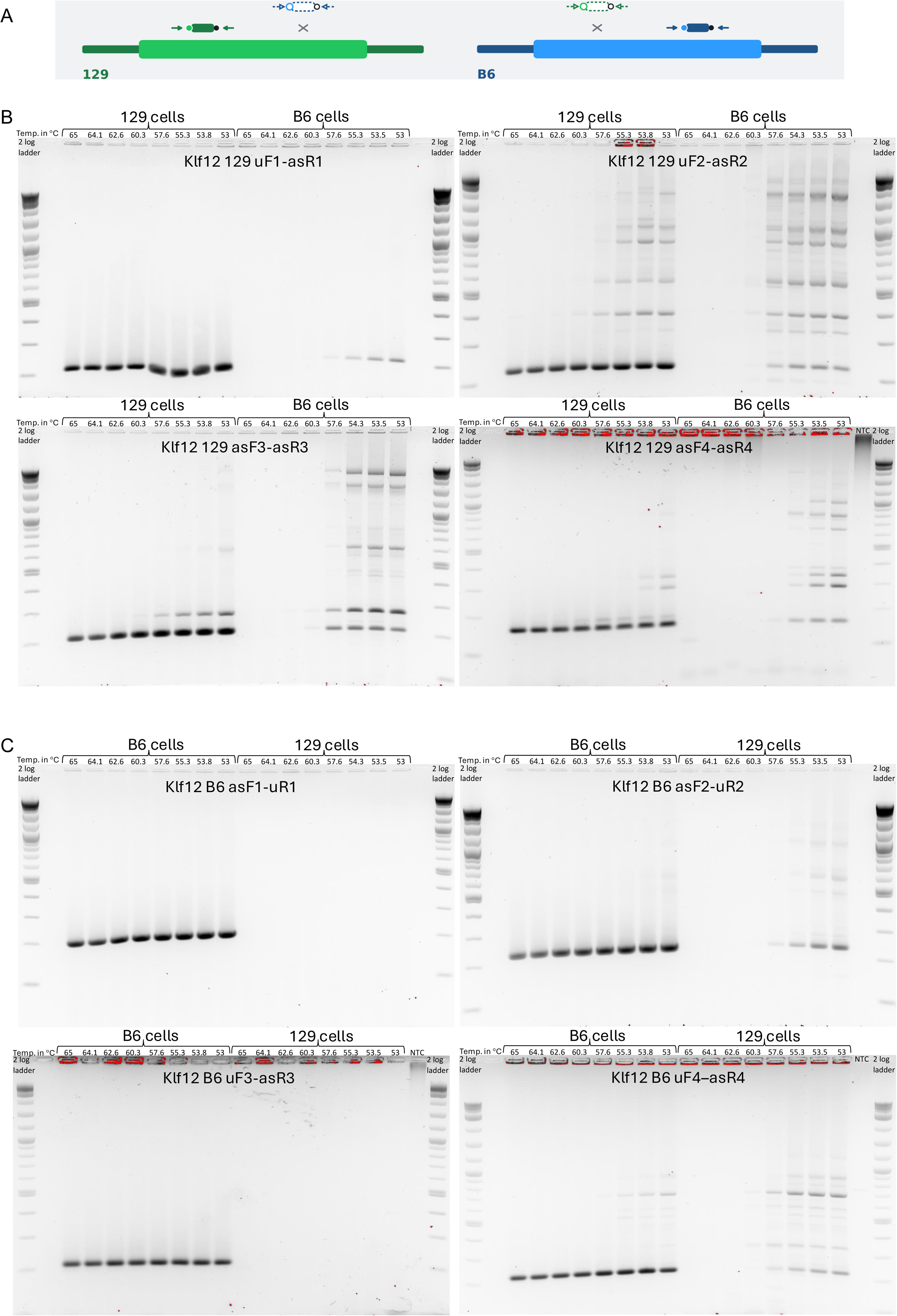
Testing allele specificity of PCR primers at *Klf12*. **A.** Schematic depicting allele specificity of primers at *Klf12*. **B.** PCR reactions with 129 allele specific primers for SNPs at *Klf12* show specificity for 129 DNA at annealing temperatures ≥60.3°C. **C.** PCR reactions with B6 allele specific primers for SNPs at *Klf12* show specificity for B6 DNA at annealing temperatures ≥60.3°C.

**Figure S2:**
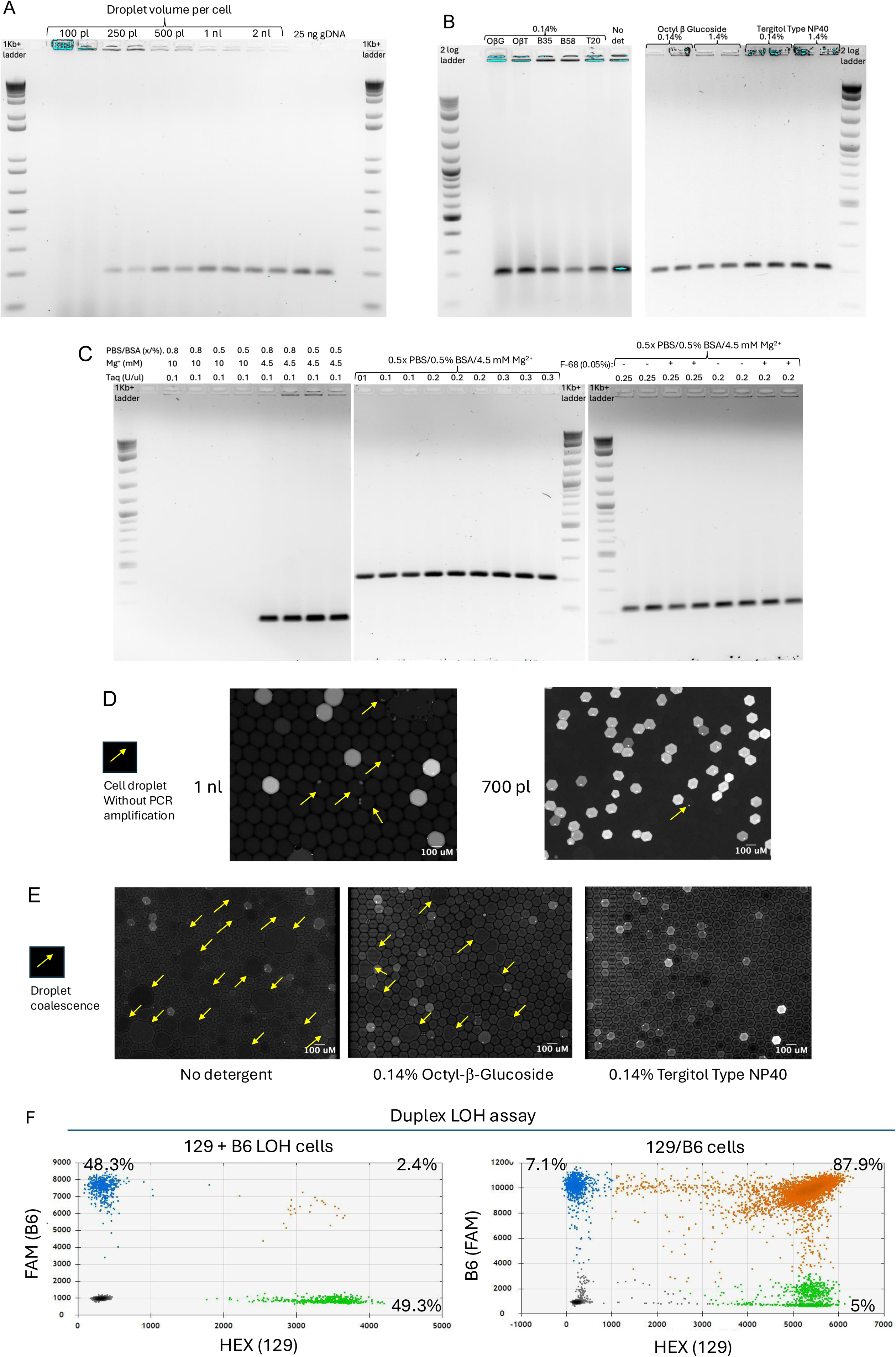
Optimization of PCR conditions for cell-in-drop genotyping using universal primers at *Klf12*. **A.** Droplet volume per cell. PCR reactions of 50 μl were tested in tubes with different numbers of cells to determine the reaction volume per single cell that gave a comparable signal compared to 25 ng genomic DNA (gDNA) in 50 μl. Therefore, 100 pl per cell indicates 500,000 cells in 50 μl PCR reaction, 200 pl indicates 250,000 cells, 500 pl indicates 100,000 cells, 1 nl indicates 50,000 cells and 2 nl indicates 25,000 cells. We found that higher cell concentrations suppressed PCR amplification: the PCR signal for cell-in-drop genotyping could be detected in reaction volumes starting at 1 cell in 250 pl and was more robust at 1 cell in 1 or 2 nl. **B.** Non-ionic detergents to stabilize emulsions during thermocycling were tested for compatibility with PCR reactions using whole cells, including octyl-β-glucoside (ObG), octyl-β-thioglucoside (ObT), Brij-35 (B35), Brij-58 (B-58), Tween 20 (T20), and Tergitol Type NP40. octyl-β-glucoside and Tergitol Type NP40 showed similar amplification as with no detergent. **C.** Different concentrations of PBS/BSA, Mg2+ and Taq and the presence of Pluronic F-68 were tested in PCR reactions with cells. Robust amplification was obtained with 0.5x PBS/0.5% BSA, 4.5 mM Mg^2+^ and 0.2 U/ml Taq. The non-ionic surfactant Pluronic F-68 (F-68; 0.05%), which can minimize cell clumping in single cell suspensions, did not affect PCR amplification. **D.** Incidences of PCR failure was frequent with cells encapsulated in ∼1nl droplets, as indicated by droplets that show a small bright focus corresponding to the cell but no fluorescence (indicated with yellow arrows). By contrast, PCR failure is very low in droplets of ∼700 pl volume. **E.** Post PCR emulsions of cells encapsulated in water-in-oil droplets were examined for droplet stability. Droplet coalescence was not observed with Tergitol Type NP40, unlike with octyl-β-glucoside or without detergent (indicated with yellow arrows). **F.** Duplexed cell-in-drop genotyping LOH assay with one set of allele specific primers/probes for each 129 (HEX) and B6 (FAM) DNA. An equal mix of 129 LOH and B6 LOH cells gives ∼2% droplets fluorescent for both genotypes, likely due to cell doublets. In unperturbed 129/B6 cells, only ∼88% of the population appeared to be heterozygous, likely due to PCR failure in the remaining cells for either the 129 or B6 allele.

**Figure S3:**
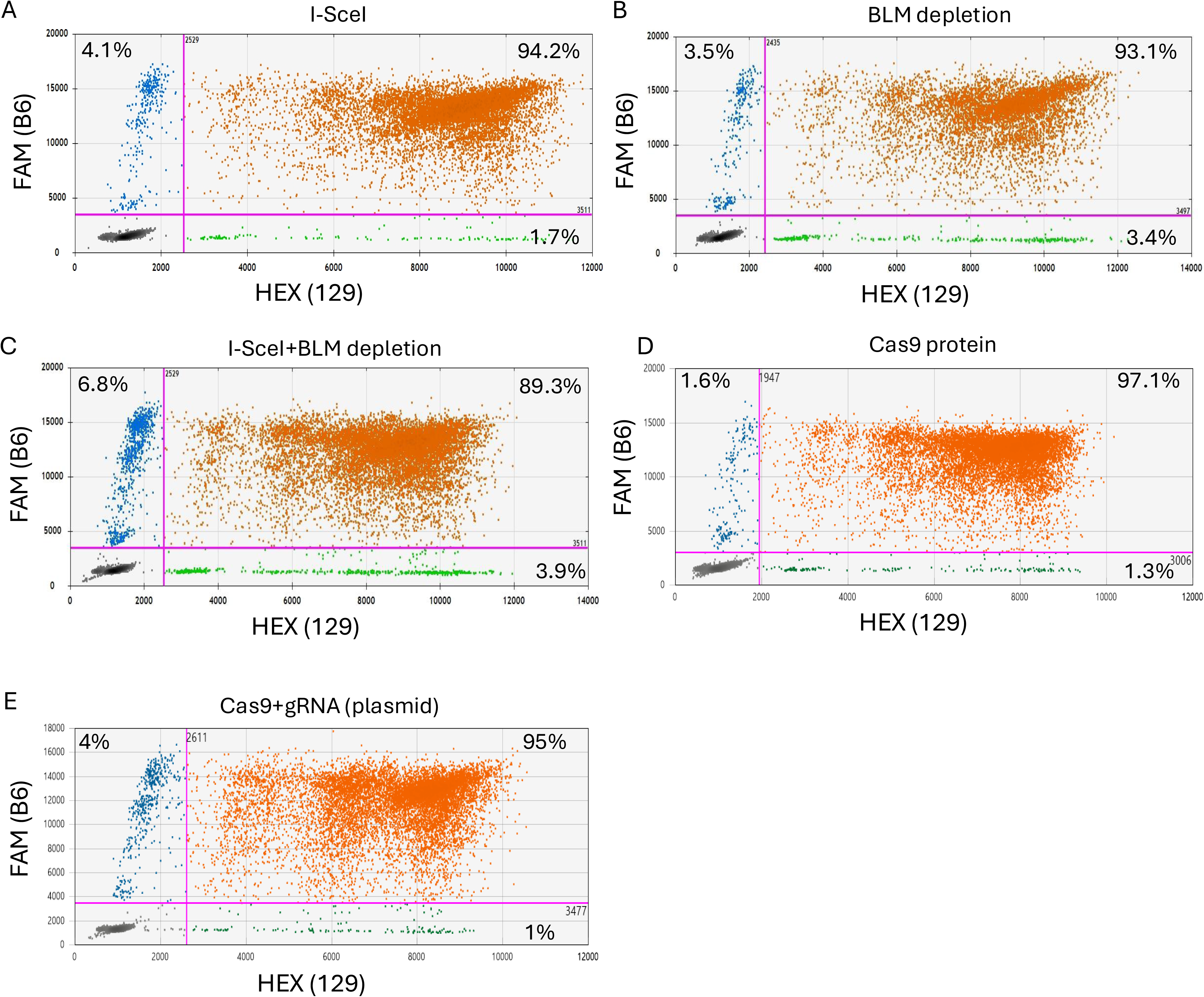
High throughput analysis for LOH at *Klf12* in cell droplets with a two-color droplet reader under various conditions. **A.E.** LOH assayed in 129/B6 cells after transfection of I-SceI expression plasmid **(A)**, BLM depletion using doxycycline **(B),** I-SceI expression and BLM depletion **(C),** transfection of Cas9 protein without gRNA (Cas9p) **(D)**, or transfection of a Cas9 and gRNA expression plasmid **(E)**. The average percent of cells in droplets from all experiments, falling within each positive quadrant, is indicated.

**Figure S4:**
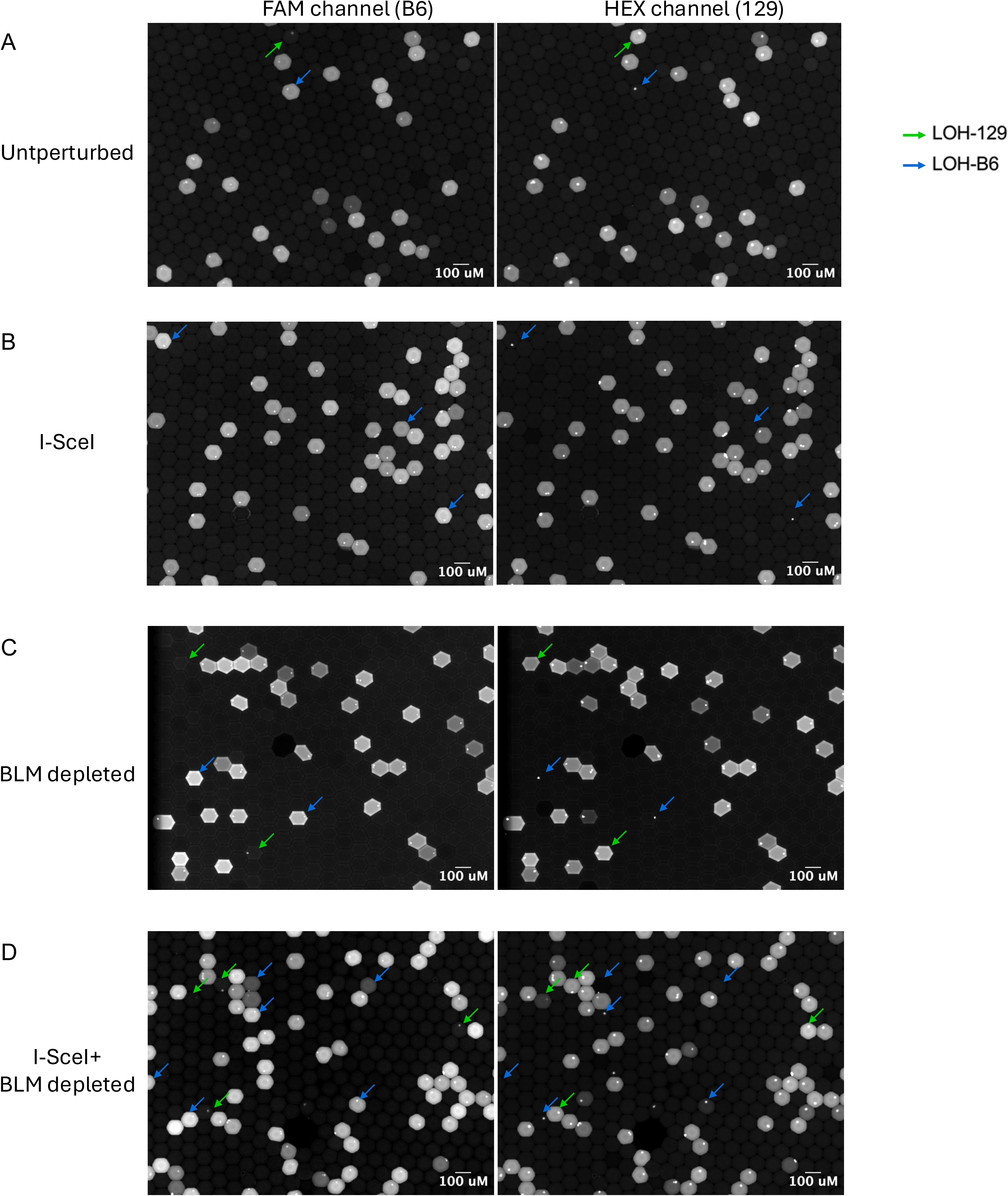
Validating high-throughput counts of multiplexed cell-in-drop genotyping droplets via manual counting of fluorescence microscopy images. **A-D.** Representative image fields showing signal with a blue laser for the FAM probes (B6 allele) and a green laser for the HEX probes (129 allele). Blue arrows indicate cell droplets inferred to have undergone B6 LOH and green arrows to have undergone 129 LOH in unperturbed cells **(A)**, cells transfected with an I-SceI endonuclease expression plasmid **(B)**, cells treated with doxycycline to deplete BLM protein **(C),** and cells transfected with an I-SceI plasmid and simultaneously treated with doxycycline to deplete BLM protein **(D).**

**Figure S5:**
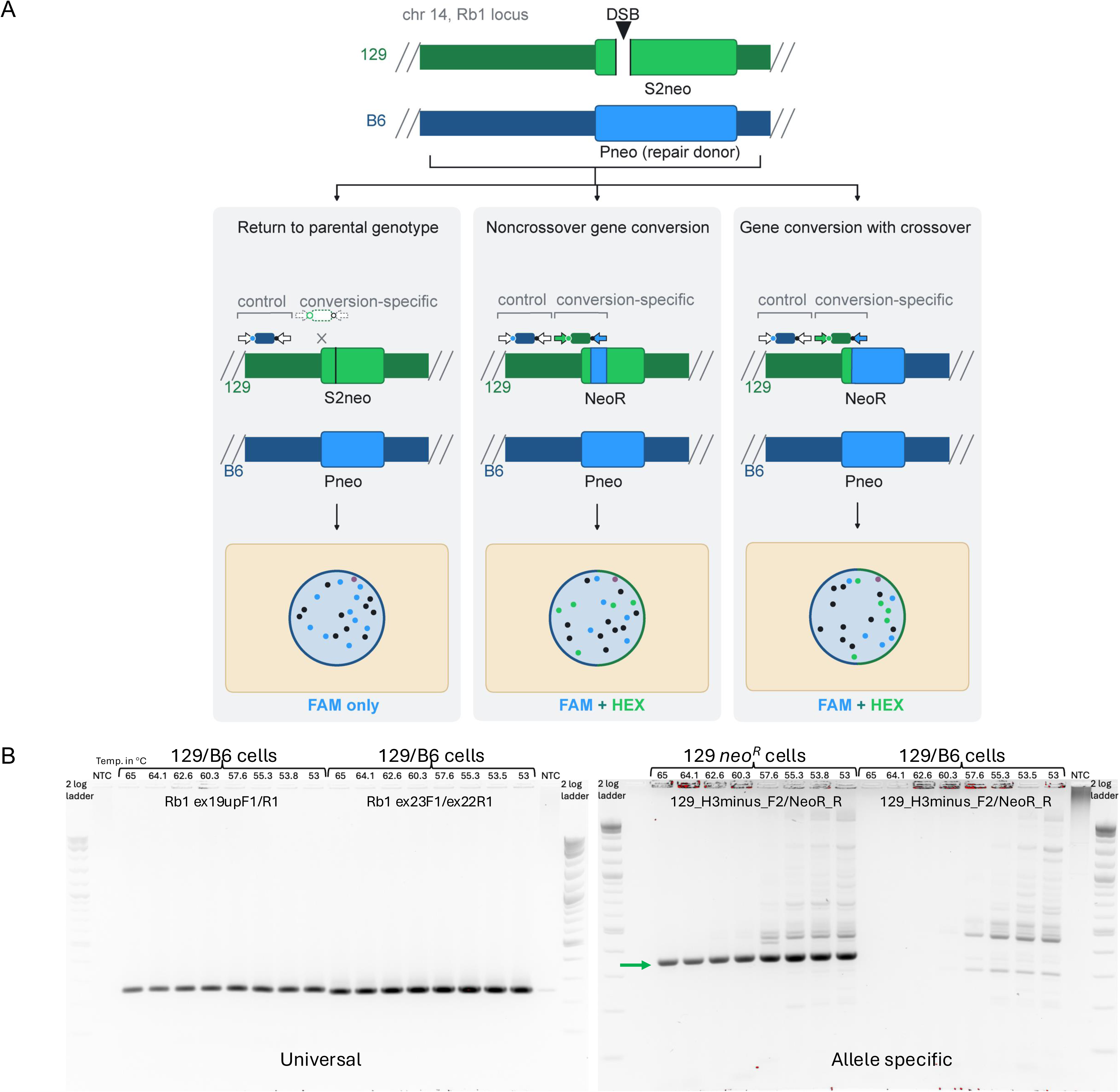
Detecting short range LOH from IH-HR using cell-in-drop genotyping. **A.** A DSB can induce IH-HR leading to short range LOH in a noncrossover event or long range LOH after a crossover event, which requires the recombinant chromatids to segregate away from each other at the next mitosis (not shown). IH-HR likely exclusively occurs during the S/G2 cell cycle phases when sister chromatids are present, although here only one chromatid is shown for simplicity. Universal PCR with FAM probes detects all cells, whereas the allele-specific PCR with a HEX probe specifically detects IH-HR events. Post PCR droplets therefore fluoresce with FAM only for non-repair events and FAM + HEX for IH-HR events. **B.** Testing PCR primers for IH-HR detection. Two sets of universal primers amplify DNA well at all tested annealing temperatures. With the allele-specific primers to detect IH-HR, specific amplification is only seen when 129 *neo^R^* LOH cells (selected from a prior IH-HR experiment) are used (green arrow), but not with the parental 129/B6 cells that are not *neo^R^*. Annealing temperatures ≥60.3°C eliminate non-specific bands.

**Figure S6:**
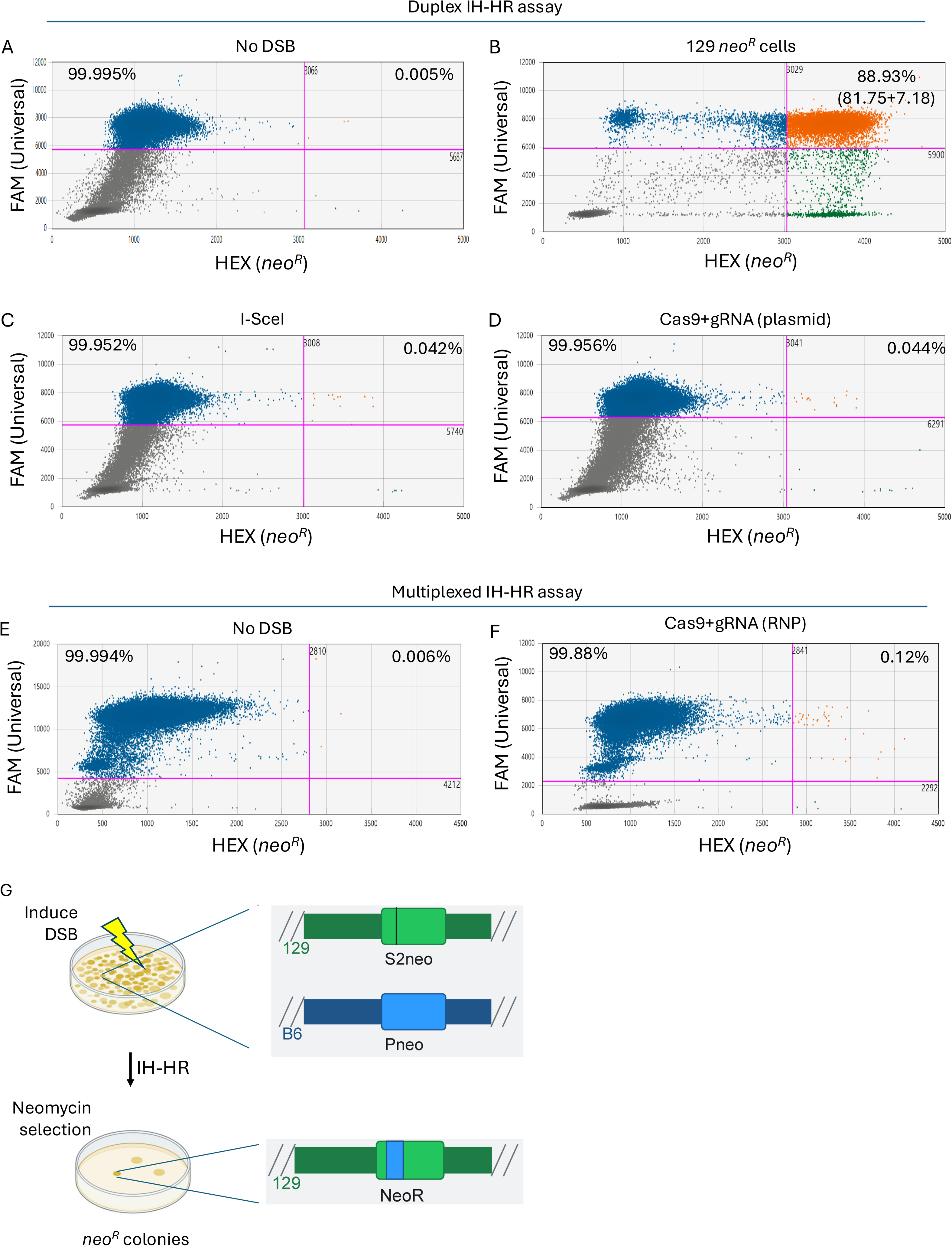
Short range LOH from IH-HR detection by cell-in-drop genotyping under various conditions **A.** Duplexed cell-in-drop genotyping IH-HR assay with one set each of universal and IH-HR primers/probes. Unperturbed heterozygous 129/B6 cells show a low background of HEX positive cell droplets (0.005%). **B.** Previously isolated 129 *neo^R^* LOH clone shows nearly 89% HEX positive droplets, ∼82% double positive and ∼7% HEX single positive. The lack of FAM signal in the latter cells may be due to allelic dropout, leading us to develop a multiplex approach with two sets universal primers/probes. Given that the cells were selected for neomycin resistance, 100% of cells should be HEX positive, rather than 89%, but the *neo^R^* gene may be difficult to amplify due to the high GC content. **C,D.** Cell-in-drop duplex assay for IH-HR in cells with a DSB induced after transfection of an I-SceI expression vector **(C)** or a Cas9 and gRNA expression vector **(D)**. The average percent of cells in droplets from all experiments, falling within each positive quadrant, is indicated. **E,F.** Cell-in-drop multiplex assay for IH-HR in cells without a DSB **(E)** or with a DSB induced after transfection of Cas9 and gRNA RNP **(F)**. The average percent of cells in droplets from all experiments, falling within each positive quadrant, is indicated. **G.** IH-HR frequency was also determined using *neo^R^* colony formation selecting for resistance to the neomycin analog G418. In IH-HR repair, a DSB at *S2neo* copies sequences from *Pneo* to result in short-range LOH and restoration of a functional neomycin resistance gene from the two defective parental neomycin alleles.

**Figure S7.**
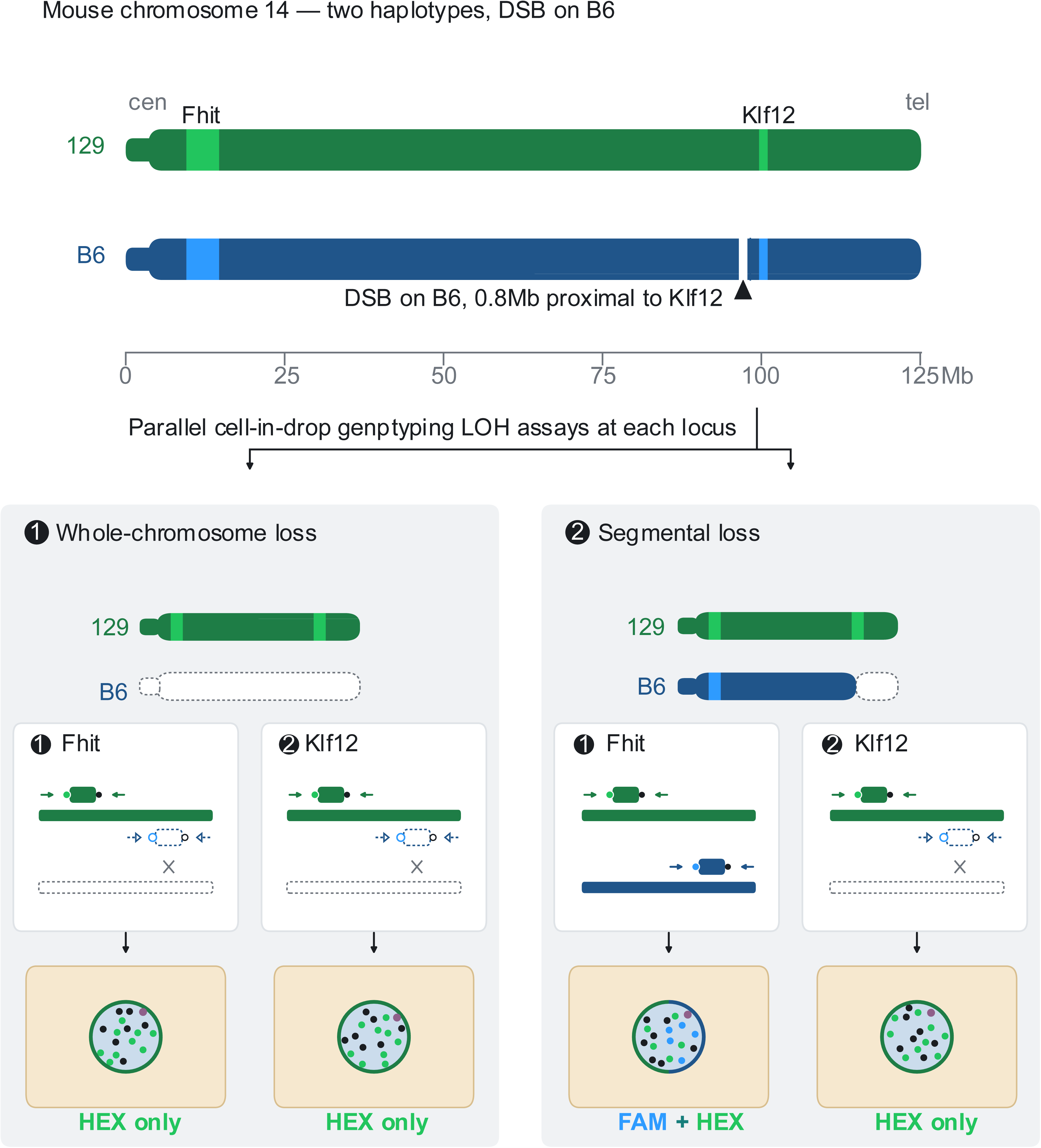
Differentiating LOH mechanisms from parallel cell-in-drop genotyping assays. Parallel cell-in-drop genotyping at centromere (*Fhit*) and telomere (*Klf12)* proximal loci in a cell population after a DSB is induced 0.8 Mb upstream of the *Klf12* locus on the B6 chromosome. Whole chromosome losses result in equivalent LOH levels at the two loci, while segmental losses lead to LOH at the telomere with retention of heterozygosity near the centromere. Post PCR droplets therefore fluoresce with HEX only at both *Fhit* and *Klf12* for cells with DSB induced whole chromosome loss but FAM + HEX at *Fhit* and HEX only at *Klf12* for DSB induced segmental loss. The DSB site for Cas9 is present on the B6 chr14 and is located 0.8 Mb upstream of *Klf12* gene in 129/B6 mESCs.

**Figure S8:**
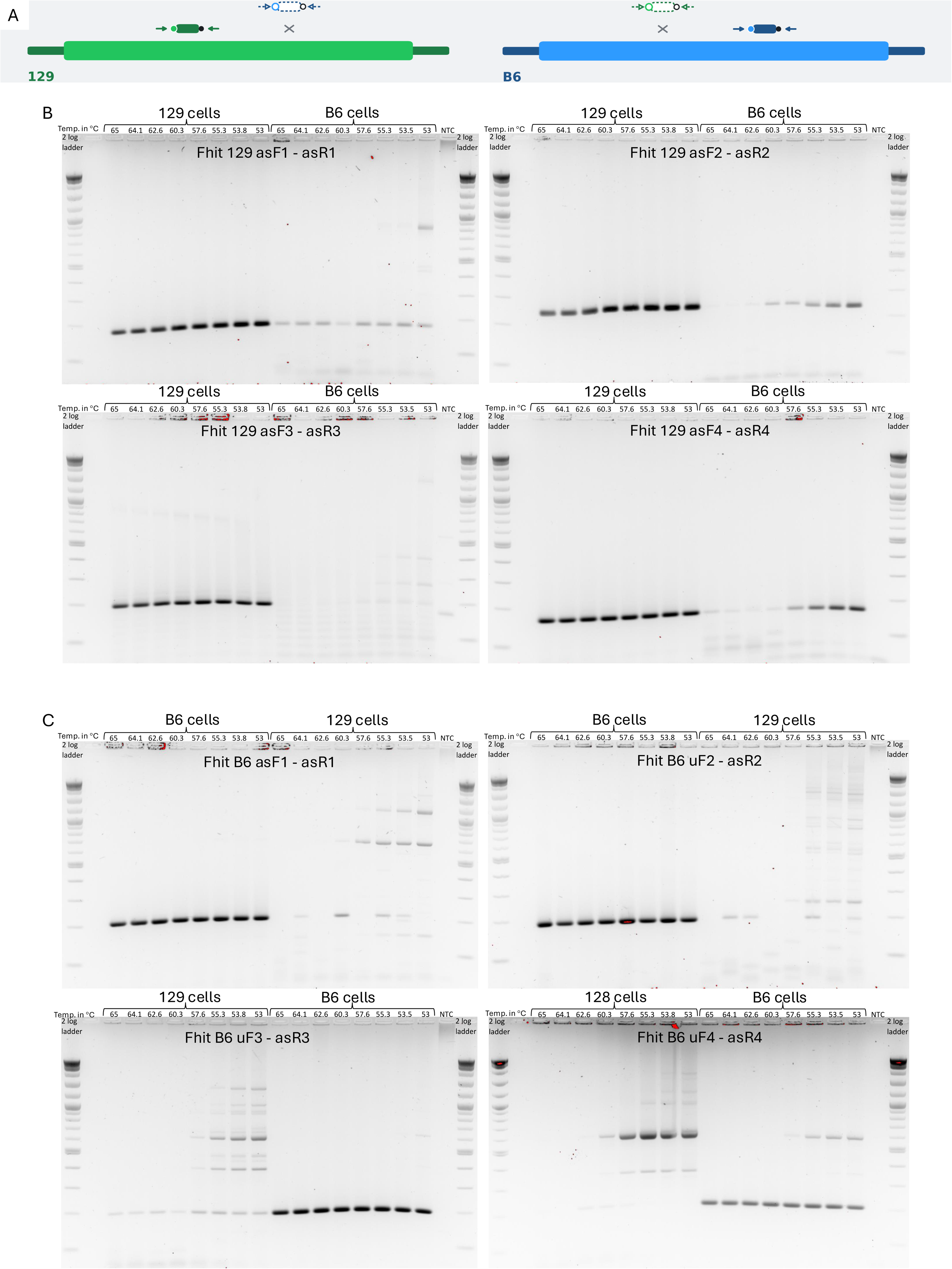
Testing allele specificity of PCR primers at *Fhit*. **A.** Schematic depicting allele specificity of primers at *Fhit*. **B.** PCR reactions with 129 allele specific primers for SNPs at *Fhit* show specificity for 129 DNA at annealing temperatures ≥60.3°C. **C.** PCR reactions with B6 allele specific primers for SNPs at *Fhit* show specificity for B6 DNA at annealing temperatures ≥60.3°C.

**Figure S9:**
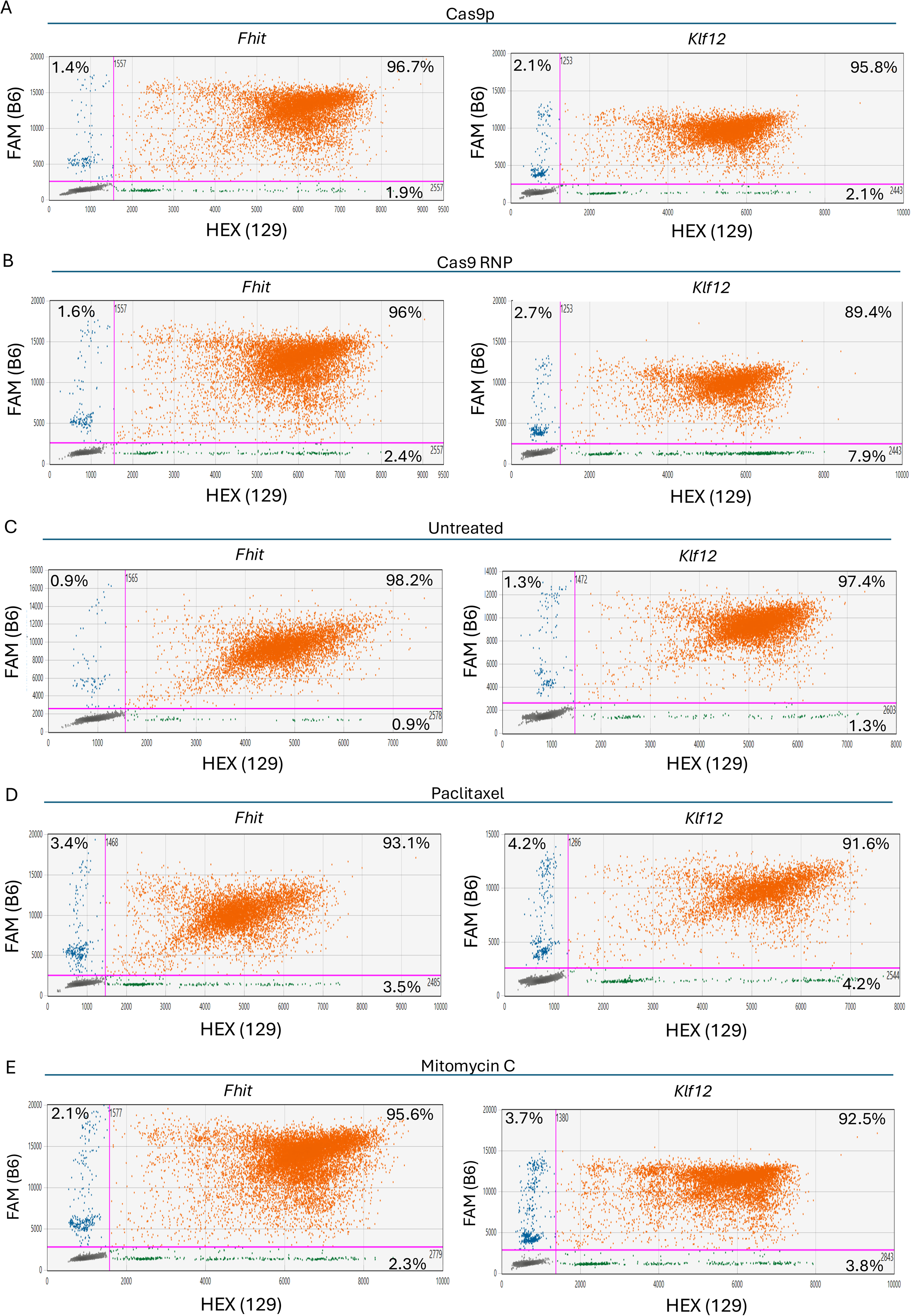
Parallel high throughput analysis for LOH at *Fhit* and *Klf12* in cell droplets with a two-color droplet reader under various conditions. **A.-E.** High throughput LOH analysis of cell droplets after transfection of Cas9 protein without gRNA (Cas9p) **(A),** transfection of Cas9 with gRNA in an RNP (Cas9 RNP) **(B)**, or of untreated cells **(C)**, Paclitaxel-treated cells **(D)**, or Mitomycin C-treated cells **(E)**. The average percent of cells in droplets from all experiments, falling within each positive quadrant, is indicated.

## Methods

### Cell culture

The 129/B6 mouse embryonic stem cells (mESCs), containing one set of chromosomes from 129 mice and one set of chromosomes from B6 mice, were previously modified to contain a tet-repressible *Blm* promoter [43] and an interhomologue homologous recombination (IH-HR) reporter [36]. mESCs were cultured in Dulbecco’s Modified Eagle Medium (Memorial Sloan Kettering Cancer Center Media Preparation Core Facility) supplemented with 12.6% Stasis stem cell qualified FBS (GeminiBio), 1% Non-Essential Amino Acids (Gibco), 1% L-glutamine (Memorial Sloan Kettering Cancer Center Media Preparation Core Facility), 1% Penicillin-Streptomycin (Gibco), 0.008% murine leukemia inhibitory factor (GeminiBio), and 0.0007% beta-mercaptoethanol (Sigma) on gelatin-coated (Memorial Sloan Kettering Cancer Center Media Preparation Core Facility) tissue culture-treated sterile plates.

### Oligonucleotide sequences

All primer, probe and crRNA sequences are listed in Tables S1 and S2.

### Transfection of mESC cells for LOH analysis

DSBs were induced via plasmid transfection at the I-SceI site at the *S2neo* gene, located at position 73.5 Mb on the 129 chr14, 3 x 10^6^ mESCs were transfected with 10 μg of plasmid expressing I-SceI [58] (pCBASceI, Addgene #26477) or Cas9-gRNA. The Cas9-gRNA expression plasmid, termed pX459+gS2Neo, was derived from cloning the gS2Neo oligo into the pSpCas9(BB)-2A-Puro plasmid (pX459) at the BbsI restriction enzyme site. For DSB induction using Cas9 RNPs, 2 x 10^6^ mESCs were transfected with *in vitro* reconstituted RNP made by mixing 140 pmol trRNA, 140 pmol crRNA, and 120 pmol Cas9 protein. The trRNA and all crRNAs (Table S1) were ordered from Integrated DNA Technologies and *S. pyogenes* Cas9 protein with an NLS was obtained from QB3 MacroLab (UC Berkeley). All nucleofections were performed using the Lonza Nucleofector 2b, program A-013, in 100 μl HM buffer in 0.2 cm cuvettes. Following transfection, all mESCs were plated on a 10-cm plate (Fisher Scientific), and then moved after 24 hours to fresh media in 6-cm plates for ∼20 hours for cells to recover before cell-in-drop genotyping.

### Genetic depletion of BLM via doxycycline treatment and chemical perturbations via Paclitaxel or Mitomycin C treatment in mESCs for cell-in-drop genotyping LOH assay

To determine the effect of BLM depletion on LOH, 1 x 10^5^ cells were plated in a 6-cm plate. The following day, cell culture media is replaced with media containing 1 µg/ml doxycycline to deplete BLM. mESCs were passaged for in media with doxycycline for 72 hours and then moved to fresh media without doxycycline for ∼20 hours for cells to recover before cell-in-drop genotyping. For combined analysis of BLM depleted and I-SceI transfected cells, 5 x 10^6^ cells were plated in a 10-cm plate with 1 µg/ml doxycycline, harvested after 24 hours to transfect with I-SceI expressing plasmid, and then replated in media with 1 µg/ml doxycycline for another 48 hours before transferring to fresh media without doxycycline in 6-cm plates for recovery as before.

To determine the effect of Paclitaxel or Mitomycin C on LOH, 1 x 10^5^ cells were plated in all wells of a 6-well plate. The following day, cell culture media is replaced with media containing 3.125nM Paclitaxel or 15.625nM Mitomycin C. mESCs are passaged in media with Paclitaxel or Mitomycin C for 72 hours and then moved to fresh media in 6-cm plates without drugs for ∼20 hours for cells to recover before cell-in-drop genotyping.

### Cell-in-drop genotyping

In order to generate single cell suspensions from adherent mESCs, 3 x 10^6^ cells mESCs are plated 24 hours post transfection or after 72 hours of doxycycline, Paclitaxel or Mitomycin C treatment in fresh media without drugs in 6-cm plates one day prior to the cell-in-drop genotyping assay and then cells are harvested ∼20 hours later. On the day of the experiment, cells were trypsinized with 0.5 ml recombinant trypsin (Thermo Fisher Scientific #12605028), incubated at 37 °C for 5 minutes, then neutralized with an equal volume of media, and trypsinized cells were made up to 3 ml by adding 1x PBS and overlaid on a 1.15 g/ml density medium consisting of Optiprep^TM^ solution in 1x PBS (Sigma-Aldrich #D1556-250ML). mESC were harvested by centrifugation at 800 rcf for 15mins, and then cells removed from the interphase layer and transferred to 200 ul of PBFO (0.5x PBS, 0.5% BSA (Non-acetylated Bovine Serum Albumin, lyophilized powder Sigma-Aldrich # B6917-5X100MG), 0.05% Pluronic™ F-68 Non-ionic Surfactant (Thermo Fisher Scientific # 24040032) and 0.15x Optiprep^TM^). Cell density was determined with an automated cell counter and cell-suspension buffer at PBFO is prepared at 1.4 cells/nl density for encapsulation.

Lysis + PCR (LP) reaction buffer for cell encapsulation was prepared by mixing 1x Jeffreys buffer (10x: 450 mM Tris-Cl pH 8.8, 110 mM (NH_4_)_2_SO_4_, 45 mM MgCl_2_, 67 mM 2-mercaptoethanol, 44 mM EDTA, 10 mM dNTPs (Roche), 1.13 mg/ml ultra-pure non-acetylated BSA (Thermo Fisher Scientific # AM2618), 12.5 mM Tris base) with 1.4% TERGITOL™ Type NP-40 solution (Sigma-Aldrich # NP40S-100ML) and 0.2U/ul Platinum™ II Taq Hot-Start DNA Polymerase (Thermo Fisher Scientific #14966005). This buffer was combined with primers and dual labeled TaqMan probes (from IDT, see table S1) for the LOH assays at *Fhit* or *Klf12* or IH-HR assay at *Rb1-neo* depending on the experiment.

LP reaction buffer and droplet oil (Qx200™ Droplet Generation Oil for EvaGreen, Bio-rad #1864005) were loaded in 1ml BD Luer-Lok^TM^ syringes (Fisher Scientific # 14-823-30), while cell-suspension was loaded in a 1 mL BD Tuberculin Syringe (Fisher Scientific # 14-823-2E). All syringes were capped with a sterile and disposable 26 ga x 1/2 inch hypodermic needle (Exel International # 26402) and medical grade polyethylene micro tubing, Dimensions: I.D. x O.D.: 0.015″ x 0.043″ / 0.38 mm x 1.09 mm (Scientific Commodities Incorporated # BB31695-PE/2). All syringes were placed in Syringe pumps (Harvard Apparatus #702226, or alternatives from, e.g., Cetoni and KD Scientific) and microtube inputs and output were loaded onto a Custom microfluidic chip (Atrandi Biosciences # MCN-Q1Q). All inputs were initially run at 500 ul/hr for inputs to stabilize the formation of water-in-oil droplets. Input rates were then adjusted to 50 ul/hr for cells, 450 ul/hr (1 in 10 dilution) for LP buffer, and ∼800 ul/hr for carrier oil, and emulsion collected after 1min 30sec of initial flow by placing the output tube in a 1.5ml Eppendorf tube on ice to collect emulsion. A 2 second video is taken at maximum frame rate and droplet volume (*V*) calculated in picoliters (pl) using the formula *V* = *Ft*/*c*_1_*N*, where F is the sum flow rate (microliters per hour) of all aqueous phases, *t* is time to count *N* number of droplet, and *c_1_* is a correction factor for unit differences, equal to 3.6 sec.ul/pl.hr [59]. Emulsions were collected ∼15 mins for LOH assays. IH-HR assays require more cells; hence emulsions were collected for ∼36 mins for those experiments.

Emulsions were transferred to wells of Eppendorf twin.tec® PCR semi-skirted, 250 µL, 96-well PCR plate (Fisher Scientific # E951020362) and sealed with pierceable foil PCR Plate Heat Seal (Bio-Rad #1814040) and thermocycled with the following conditions: initial denaturation at 95°C for 20 minutes, followed by 45 cycles of denaturation at 90°C for 1 minute and annealing + extension at 60.3°C for 2 minutes, followed by resting at 7°C. All steps have a 1°C/sec ramp rate. Post PCR droplets can be loaded onto cell counting chamber slides (Thermo Fisher Scientific #C10312) and manually examined under fluorescent microscopy or analyzed at high throughput with the Bio-Rad QX200 digital droplet PCR reader (Bio-Rad #1864001).

